# SNX2, but not SNX1, regulates the hepatocyte growth factor receptor MET in HCT116 colorectal cancer cells

**DOI:** 10.64898/2026.10.06.755412

**Authors:** Laiyen Garcia Delgado, Amélie Derome, Christine L. Lavoie, Caroline Saucier, Jean-Bernard Denault

**Author notes:** Correspondence: Jean-Bernard Denault. These authors should be considered joint first authors (Laiyen Garcia Delgado, Amélie Derome). These authors should be considered joint senior authors (Christine L. Lavoie, Caroline Saucier, Jean-Bernard Denault).

## Abstract

Aberrant signaling of receptor tyrosine kinases (RTKs), including the hepatocyte growth factor receptor MET, contributes to colorectal cancer (CRC) progression and is tightly regulated by endocytic trafficking. We previously showed that combined deletion of the endosomal sorting proteins SNX1 and SNX2 disrupts MET trafficking and enhances downstream signaling in HCT116 CRC cells. However, their individual contributions remained unresolved. Using CRISPR/Cas9-engineered *SNX1*-and *SNX2*-KO HCT116 cells, we examined their roles in MET trafficking and signaling.

Loss of *SNX2*, but not *SNX1*, markedly reduced SNX5 and SNX6 protein levels without affecting transcript abundance. *SNX2* deficiency also increased total MET under serum-containing conditions without altering *MET* transcript or cell-surface receptor levels. Following HGF stimulation, loss of *SNX2*, but not *SNX1*, transiently delayed MET trafficking to EEA1^+^ early and RAB7^+^ late endosomes, while recycling and lysosomal trafficking remained largely unaffected. This was accompanied by enhanced and sustained HGF-induced AKT activation.

Together, these findings identify SNX2 as a predominant regulator of early MET endosomal trafficking in HCT116 cells and reveal functional asymmetry between SNX1 and SNX2. They support a role for SNX2 in spatial regulation of MET signaling through receptor trafficking, consistent with the association of low SNX2 expression with poorer survival in CRC patients.

## INTRODUCTION

Spatial regulation and location-biased signaling have emerged as crucial mechanisms governing receptor tyrosine kinase (RTK) activity [1,2]. Rather than signaling exclusively from the plasma membrane following ligand stimulation, internalized RTKs can assemble signaling complexes as they transit through the various endosomal compartments [3–5]. For instance, while initial activation of RTKs at the cell surface triggers localized RAS-ERK signaling, receptor retention in early endosomes (EEs) is frequently required to sustain prolonged PI3K-AKT activation [3,4]. Given the critical role of dysregulated RTK signaling in diseases such as cancer, understanding how alterations in receptor trafficking and intracellular localization shape signaling outputs and cellular responses is therefore important.

After internalization, often via clathrin-mediated endocytosis, RTKs cluster in EEs (sorting endosomes), where they are routed either to lysosomes for degradation or back to the plasma membrane via recycling pathways. Sorting proteins recruited to EEs regulate the intracellular fate of activated receptors, including members of the sorting nexin (SNX) family. Members of this family are characterized by a phox-homology (PX) domain that binds membrane phospholipids enriched in endosomal compartments [6]. A subgroup of SNXs, called SNX-BAR (including SNX1, SNX2, SNX5, and SNX6), also contains an additional C-terminal tubule-forming BAR (Bin/Amphiphysin/Rvs) domain [7]. To function as EE sorting factors, SNX1 and SNX2 preferentially heterodimerize with SNX5 and SNX6 to form the ESCPE-1 complex (Endosomal SNX-BAR Sorting Complex for Promoting Exit 1), driving endosomal cargo exit and retrograde trafficking [8–11].

SNX-BAR proteins are implicated in RTK intracellular trafficking [12–15]. Although SNX1 and SNX2 share high sequence identity and have historically been viewed as largely redundant components of endosomal sorting machinery [12,13,16], their individual contributions to RTK trafficking and signaling remain poorly defined. However, their effects on RTK trafficking and attenuation appear highly context-dependent and vary across tissue types and genetic backgrounds. For instance, downregulation of SNX1 or SNX2 in lung cancer models accelerates MET endocytosis and drives a delayed, sustained peak in receptor phosphorylation prior to lysosomal clearance [17,18], whereas silencing *SNX2* expression in cervical carcinoma HeLa cells potentiates HGF-induced MET and ERK1/2 activation [19]. In contrast, our recent work in HCT116 colorectal cancer (CRC) cells showed that combined knockout (KO) of SNX1 and SNX2 (*SNX1/2*-dKO) delays early endosomal transit of MET upon HGF stimulation, sustains higher MET and AKT activation, and enhances resistance to death ligand-induced apoptosis [20]. Although both SNX1 and SNX2 expression are reduced in CRC tumors, low SNX2, but not SNX1, expression has been associated with poorer patient survival [19,21], further supporting the importance of determining whether these trafficking and signaling defects stem from loss of SNX1, SNX2, or their combined deficiency.

In this study, we generated single CRISPR/Cas9 KO HCT116 cell lines to isolate the functional roles of SNX1 and SNX2 in regulating MET. We demonstrate that loss of SNX2, but not SNX1, leads to a co-regulated reduction in SNX5/6 protein levels, delayed MET entry into EEA1^+^ EEs, and potentiated AKT phosphorylation following HGF stimulation. These findings support a distinct role for SNX2 in regulating the spatiotemporal trafficking of MET and downstream AKT activation in HCT116 CRC cells.

## RESULTS

### SNX2, but not SNX1, Deletion Reduces SNX5 and SNX6 Protein Levels

To investigate the individual contributions of SNX1 and SNX2 to receptor trafficking, single CRISPR/Cas9 KO HCT116 cell lines were established. Genomic sequencing confirmed an 11-nucleotide deletion in both alleles of *SNX1* and a frame-shifting mutation in *SNX2*, consisting of a 3-nucleotide deletion and a 1-nucleotide insertion near the respective Cas9 target sites (Fig. S1). These alterations resulted in the complete loss of SNX1 and SNX2 protein expression, respectively (Fig. 1A). Single KO cells displayed no apparent morphological changes and grew at the same rate as the parental cells under standard culture conditions (data not shown).

**Figure 1:**
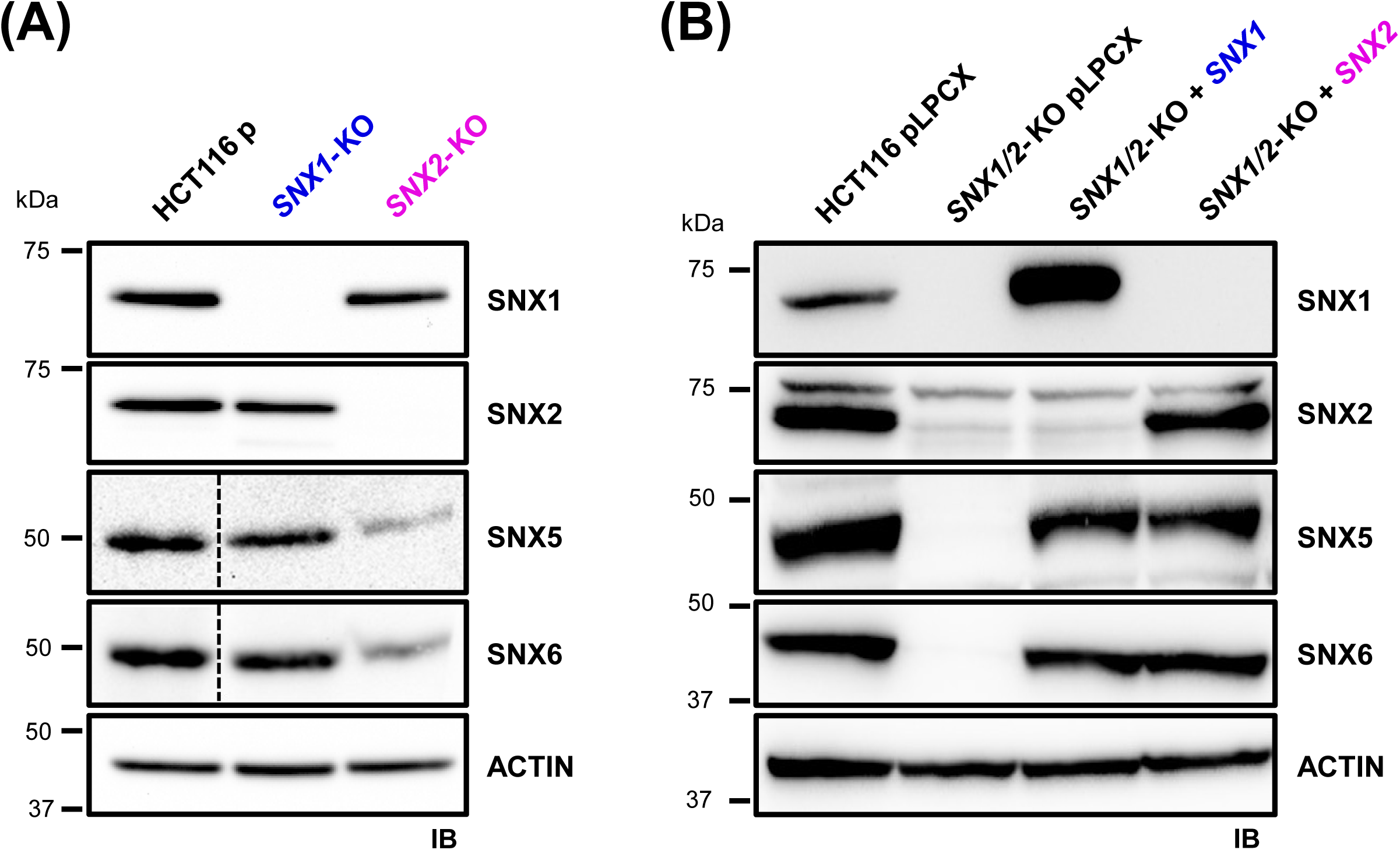
Co-regulation of SNX1, SNX2, SNX5, and SNX6 protein expression. **A)** Immunoblot analysis of steady-state SNX1, SNX2, SNX5, and SNX6 protein levels in parental (p), *SNX1*-KO, and *SNX2*-KO HCT116 cells. Dashed vertical lines indicate where non-adjacent lanes from the same gel were joined for presentation purposes. **B)** Retroviral ectopic re-expression of *SNX1* or *SNX2* in *SNX1/2-*dKO HCT116 cells restores SNX5 and SNX6 protein levels.

SNX-BAR family members assemble into higher-order multimeric complexes, with SNX1 and SNX2 specifically heterodimerizing with SNX5 and SNX6 within the ESCPE-1 [11]. In our previous work, we reported that SNX5 and SNX6 protein levels were greatly reduced in *SNX1/2*-dKO HCT116 cells [20]. Herein, we evaluated whether single deletion of *SNX1* or *SNX2* alters the levels of these partner proteins. Immunoblotting analysis of single KO lines revealed that loss of *SNX1* had no major effect on SNX5 or SNX6 protein levels (Fig. 1A). In stark contrast, loss of *SNX2* alone resulted in a marked reduction in both SNX5 and SNX6 (Fig. 1A), recapitulating the phenotype observed in dKO cells (Fig. 1B) [20]. Because *SNX5* and *SNX6* mRNA levels measured by qRT-PCR (data not shown) remained unchanged, the observed reduction in SNX5 and SNX6 protein expression in *SNX2*-KO cells suggests a post-transcriptional co-regulatory mechanism [11,22].

To determine whether the loss of SNX5 and SNX6 was reversible, we performed rescue experiments in *SNX1/2*-dKO HCT116 cells. Stable retroviral re-expression of either *SNX1* or *SNX2* in *SNX1/2*-dKO cells restored both SNX5 and SNX6 protein expression to levels comparable to parental controls (Fig. 1B). Notably, while re-expression of *SNX2* restored SNX2 protein to levels matching those in parental cells, *SNX1* re-expression resulted in much higher SNX1 protein levels than in control cells (Fig. 1B).

Taken together, these data indicate a functional asymmetry between SNX1 and SNX2 in maintaining complex stability. Endogenous SNX2 plays a primary role in maintaining SNX5 and SNX6 protein expression in HCT116 cells, whereas physiological levels of SNX1 are insufficient to prevent SNX5/6 decrease in the absence of SNX2. Therefore, the loss of SNX5 and SNX6 observed in *SNX1/2*-dKO cells is driven largely by SNX2 deficiency.

### Loss of SNX2, but Not SNX1, Promotes Intracellular Accumulation of MET Under Steady-State Growth Conditions

Having established the differential effects of SNX1 and SNX2 loss on ESCPE-1 components, we next evaluated whether the loss of *SNX1* or *SNX2* affects steady-state MET receptor expression. In *SNX2*-KO HCT116 cells cultured under standard growth conditions (10% FBS), total MET protein levels increased by 1.6-fold compared to parental cells, whereas *SNX1*-KO cells displayed MET levels comparable to parental cells, as determined by immunoblotting (Fig. 2A). Notably, this increase in MET expression was serum-dependent, as following 16 h of serum starvation, total MET levels in *SNX2*-KO cells returned to baseline levels matching parental controls (Fig. 2A).

**Figure 2:**
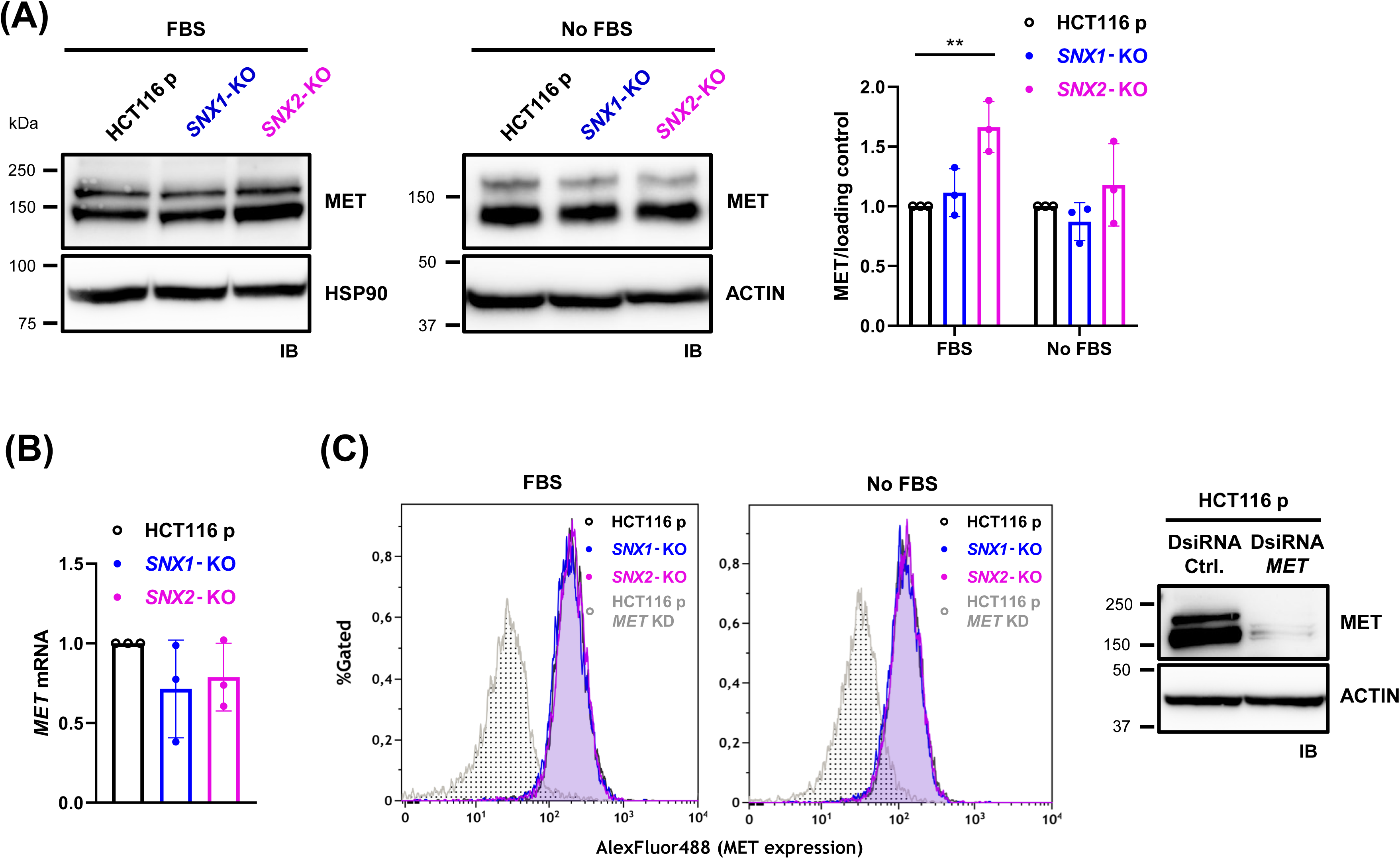
Loss of *SNX2* leads to intracellular accumulation of MET protein under serum-stimulated conditions. **A)** Immunoblot analysis of total MET protein levels in parental, *SNX1*-KO, and *SNX2*-KO HCT116 cells grown in complete medium supplemented with 10% FBS or under serum-starved conditions. Heat shock protein 90 (HSP90) and β-Actin served as loading controls. The graph shows normalized MET quantification. Data correspond to the mean ± SD (*N*=3). **B)** Quantitative RT-PCR analysis of *MET* mRNA levels across cell lines, normalized to parental levels. Data correspond to the mean ± SD (*N*=3). **C)** Cell-surface MET expression measured by flow cytometry in cells cultured in FBS-containing medium or serum-deprived conditions. DsiRNA *MET* knockdown (KD) in HCT116 cells served as a control for antibody specificity (gray trace). Knockdown efficiency was verified by immunoblotting, with β-Actin serving as a loading control.

Quantitative RT-PCR demonstrated that *MET* mRNA levels remained unchanged across parental and single-KO cell lines, ruling out transcriptional upregulation as the basis for the increase in MET protein levels (Fig. 2B). Flow-cytometry analysis of unpermeabilized cells, validated for antibody specificity using *MET*-knockdown HCT116 controls, revealed no genotype-dependent differences in cell-surface MET density among parental, *SNX1*-KO, and *SNX2*-KO cells under either serum-fed or serum-starved conditions (Fig. 2C).

Together, these results indicate that loss of *SNX2* leads to a serum-dependent increase in total MET without a corresponding increase in *MET* gene expression or cell-surface receptor density, consistent with the accumulation of an intracellular MET pool in *SNX2*-KO cells.

### Loss of SNX2, but Not SNX1, Transiently Delays MET Endosomal Trafficking

Our previous work demonstrated that MET delivery to EEA1^+^ EEs is delayed in HCT116 *SNX1/2*-dKO cells, whereas downstream routing to RAB7^+^ late endosomes, fast and slow recycling compartments (GGA3^+^ and CD71^+^, respectively), and LAMP1^+^ lysosomes remains comparable to parental cells [20]. Therefore, we next evaluated whether the individual loss of *SNX1* or *SNX2* recapitulates this early endosomal trafficking defect.

We tracked the spatial colocalization of cell-surface labeled MET with the early endosomal marker EEA1 over a 30-min period following HGF stimulation (Fig. 3). Before stimulation, MET predominantly localized to the plasma membrane in parental, *SNX1*-KO, and *SNX2*-KO cells. At 7 min after HGF stimulation, MET was readily present in EEA1^+^ endosomes in parental and *SNX1*-KO cells. In contrast, *SNX2*-KO cells displayed less MET within EEA1^+^ endosomes, with a greater proportion of the receptor remaining at or near the plasma membrane. By 15 and 30 min, MET distribution and colocalization with EEA1 appeared comparable across all three cell lines. (Fig. 3B, C). Spatial overlap between MET and EEA1 was quantified using Pearson’s correlation coefficient (PCC), with values approaching +1 indicating a strong positive correlation between fluorescence intensities in the two channels [23,24]. *SNX1*-KO cells displayed MET-EEA1 colocalization kinetics comparable to parental cells throughout the time course (Fig. 3B). In contrast, *SNX2*-KO cells showed a significantly reduced MET-EEA1 colocalization at 7 min post-stimulation (0.14 ± 0.05 *vs.* 0.31 ± 0.07 in parental cells), whereas colocalization at 15 and 30 min was comparable to the parental cell lines (Fig. 3C).

**Figure 3:**
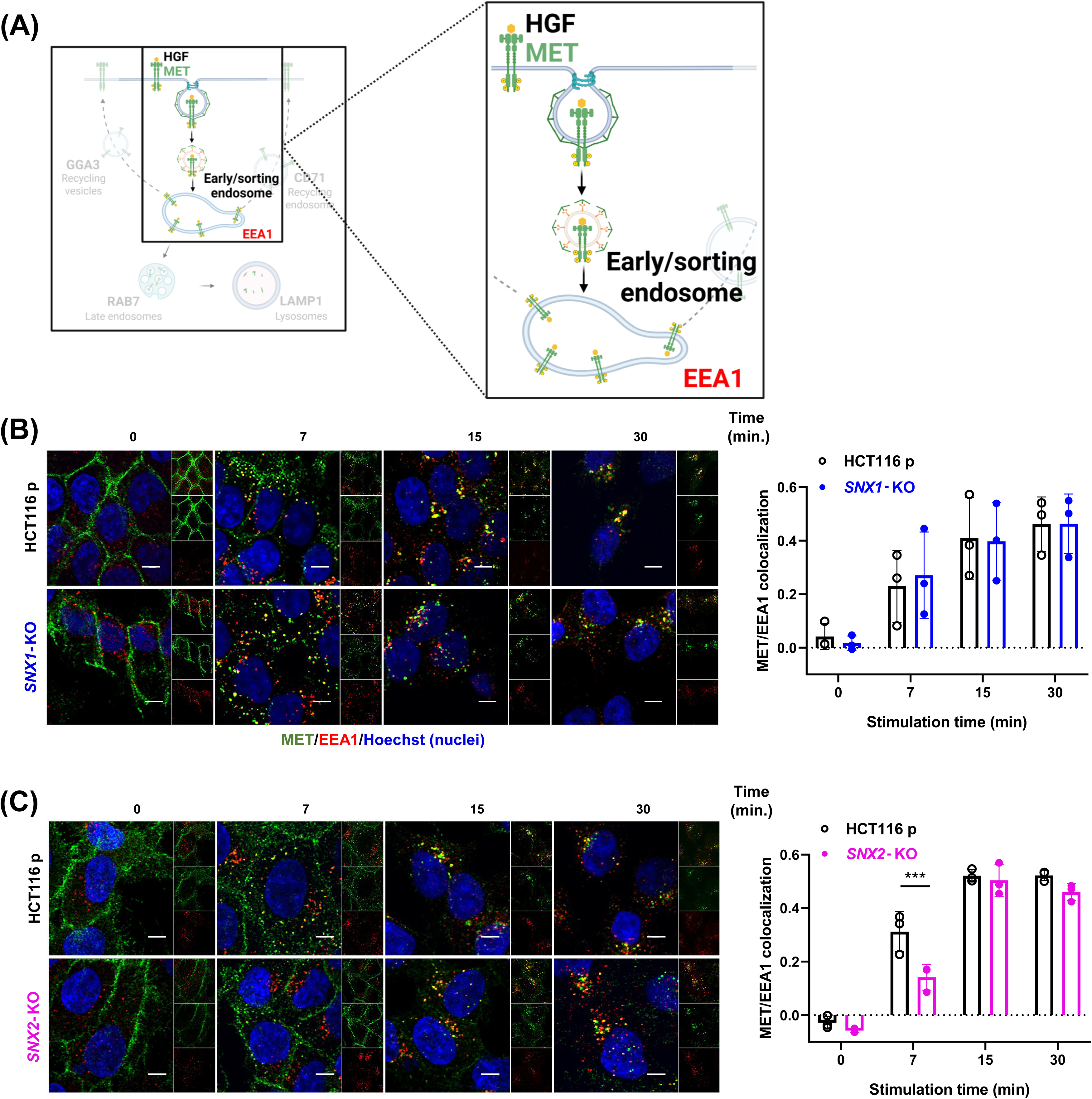
Loss of *SNX2* selectively delays MET trafficking to EEs upon HGF stimulation. **A)** Schematic representation of MET trafficking to EEA1^+^ endosomes. **B)**, **C)** Representative confocal immunofluorescence micrographs showing MET and EEA1 colocalization in parental and single *SNX1-***(B)** or *SNX2-***(C)** KO HCT116 cells. Prior to stimulation with HGF (40 ng/mL) for the indicated periods, cell-surface MET was labeled with the anti-MET L6E7 antibody (green). EEs were labeled with EEA1 (red), and nuclei were counterstained with Hoechst (blue). Scale 10 μm; 60X objective. Colocalization was quantified using the PCC. Data represent the mean ± SD (*N*=3).

We next evaluated MET transit through RAB7^+^ late endosomes following HGF stimulation (Fig. 4). Consistent with the sequential progression of internalized receptors from early to late endosomal compartments, MET colocalization with RAB7 followed the initial recruitment of MET to EEA1^+^ early endosomes. *SNX1*-KO cells displayed MET-RAB7 colocalization kinetics comparable to parental cells at all time points. In contrast, *SNX2*-KO cells exhibited a significant decrease in MET-RAB7 colocalization at 7 min post-stimulation (0.01 ± 0.04) compared with parental and *SNX1*-KO cells (0.08 ± 0.02 and 0.08 ± 0.05, respectively; Fig. 4B,C). At 15 and 30 min, MET-RAB7 colocalization in *SNX2*-KO cells was comparable to levels observed in parental cells.

**Figure 4:**
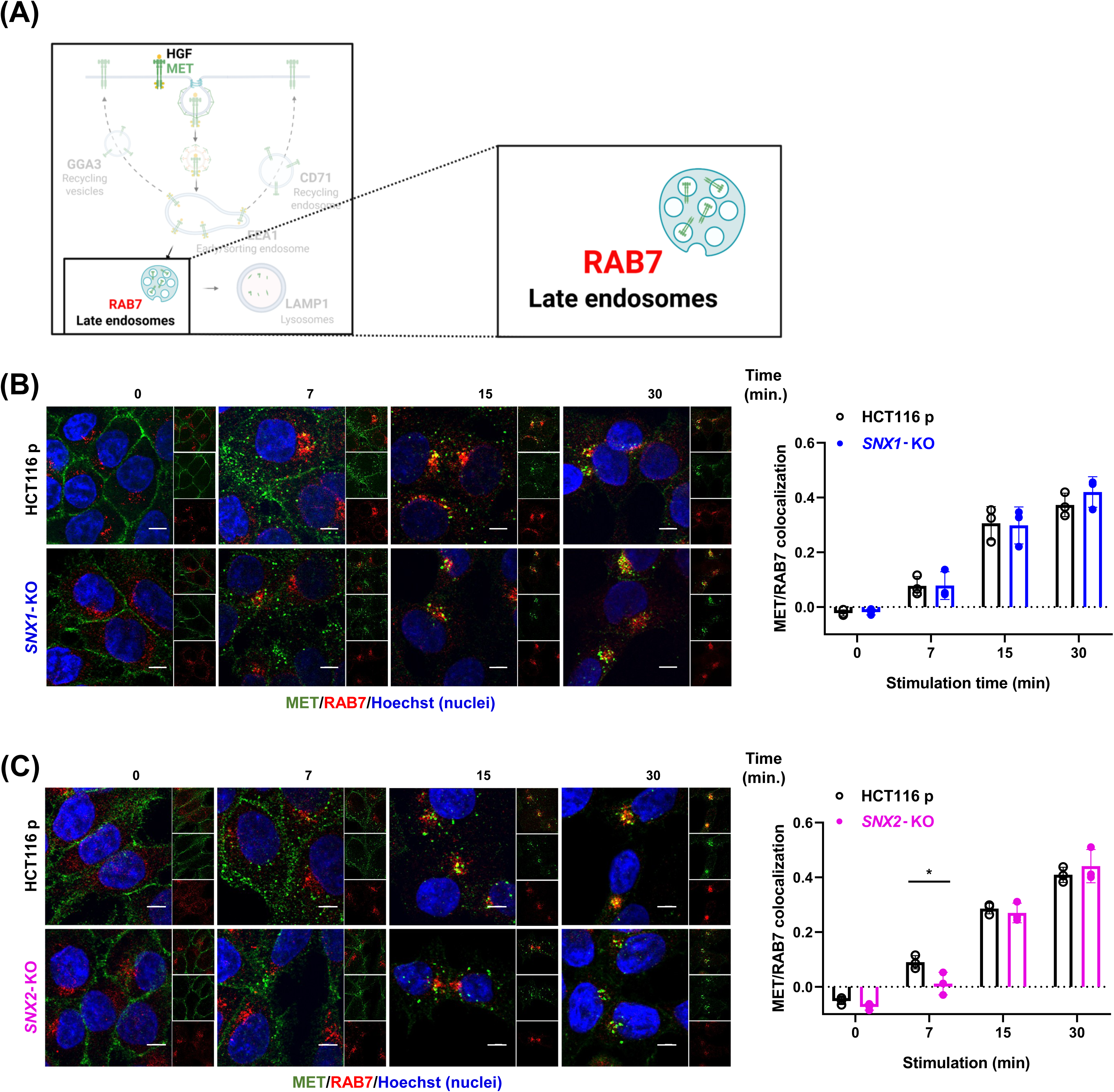
Loss of *SNX2* transiently delays MET trafficking to RAB7^+^ late endosomes. **A)** Schematic representation of MET trafficking to RAB7^+^ late endosomes. **B)**, **C)** Representative confocal immunofluorescence micrographs of the co-staining of MET and RAB7^+^ in parental and single *SNX1-***(B)** or *SNX2-***(C)** KO HCT116 cells. Prior to stimulation with HGF (40 ng/mL) for the indicated periods, surface MET was labeled with the anti-MET L6E7 antibody (green). Late endosomes were labeled for RAB7 (red), and nuclei were counterstained with Hoechst (blue). Scale 10 μm; 60X objective. Quantification of colocalization was assessed using the PCC. Data represent the mean ± SD (*N*=3).

Together, these results indicate that loss of SNX2, but not SNX1, transiently delays MET trafficking to both EEA1^+^ early endosomes and RAB7^+^ late endosomes following HGF stimulation.

### Loss of SNX1 or SNX2 Does Not Alter MET Trafficking to Lysosomal or Recycling Compartments

Because a significant portion of internalized MET is routed to lysosomes for degradation [25] we tested whether loss of *SNX1* or *SNX2* affects MET trafficking to LAMP1^+^ lysosomes. Although MET-LAMP1 colocalization increased over time following HGF stimulation, no statistically significant differences were observed between parental HCT116 cells and either single-KO cell line at any time point examined (Fig. 5).

**Figure 5:**
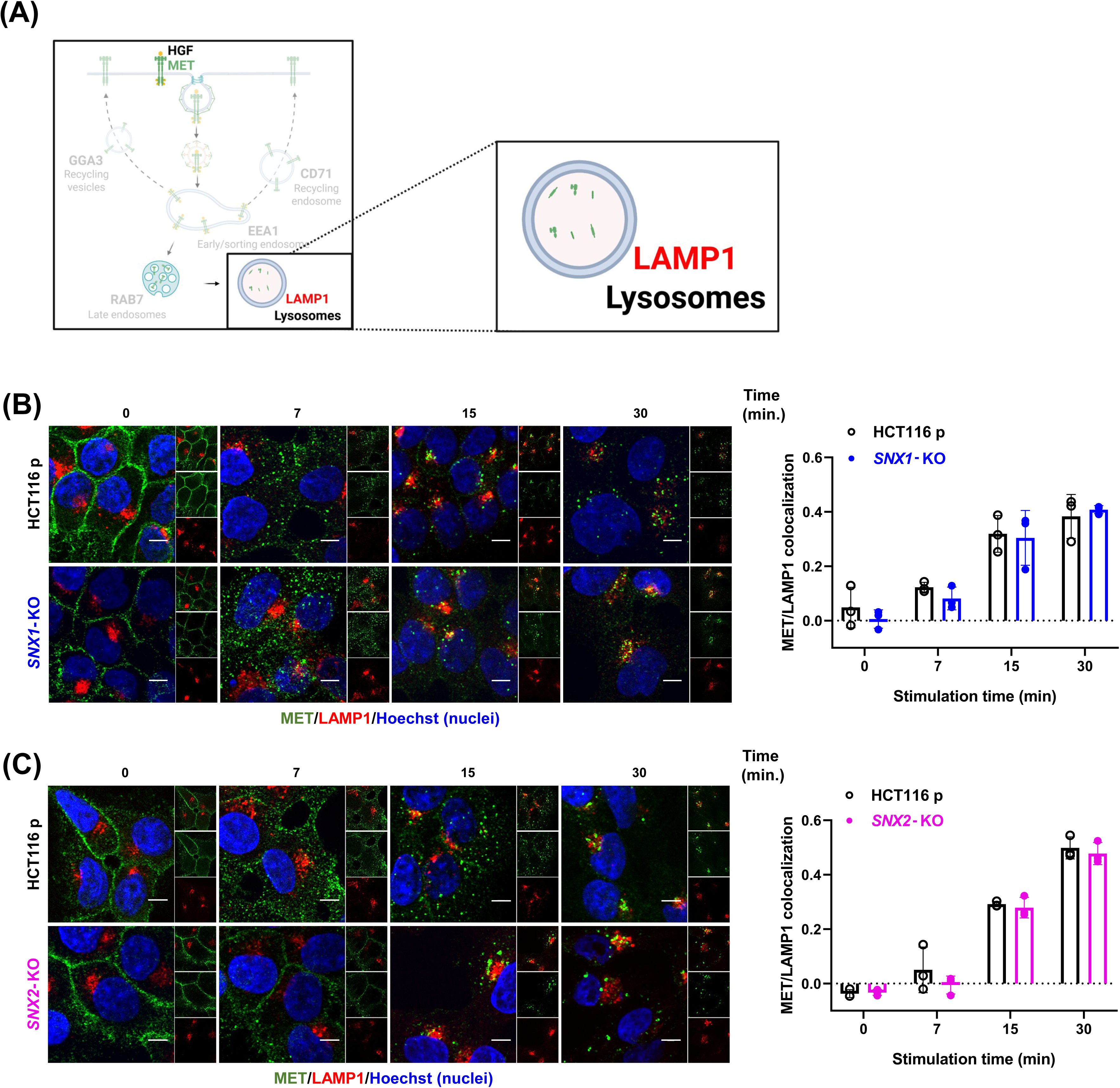
Loss of *SNX1* or *SNX2* does not alter MET trafficking to LAMP1-positive lysosomes following HGF stimulation. **A)** Schematic representation of MET trafficking to LAMP1-positive lysosomes. **B**, **C)** Representative confocal immunofluorescence micrographs showing co-staining of MET and LAMP1 in parental and single *SNX1* **(B)** or *SNX2-*KO **(C)** HCT116 cells. Prior to stimulation with HGF (40 ng/mL) for the indicated periods, cell-surface MET was labeled with the anti-MET L6E7 antibody (green). Lysosomes were labeled with anti-LAMP1 (red), and nuclei were counterstained with Hoechst (blue). Scale bar, 10 μm; 60X objective. Colocalization was quantified using PCC, as described in MATERIAL AND METHODS. Data represent the mean ± SD (*N*=3).

As an alternative to lysosomal degradation, internalized MET can be recycled back to the plasma membrane (Fig. 6A). We therefore examined whether MET trafficking through recycling compartments was affected. Similarly to our findings for LAMP1, no significant differences were observed between parental and single-KO cell lines in MET colocalization with markers of fast (GGA3; Fig. 6B) or slow recycling endosomes (CD71; Fig. 6C).

**Figure 6:**
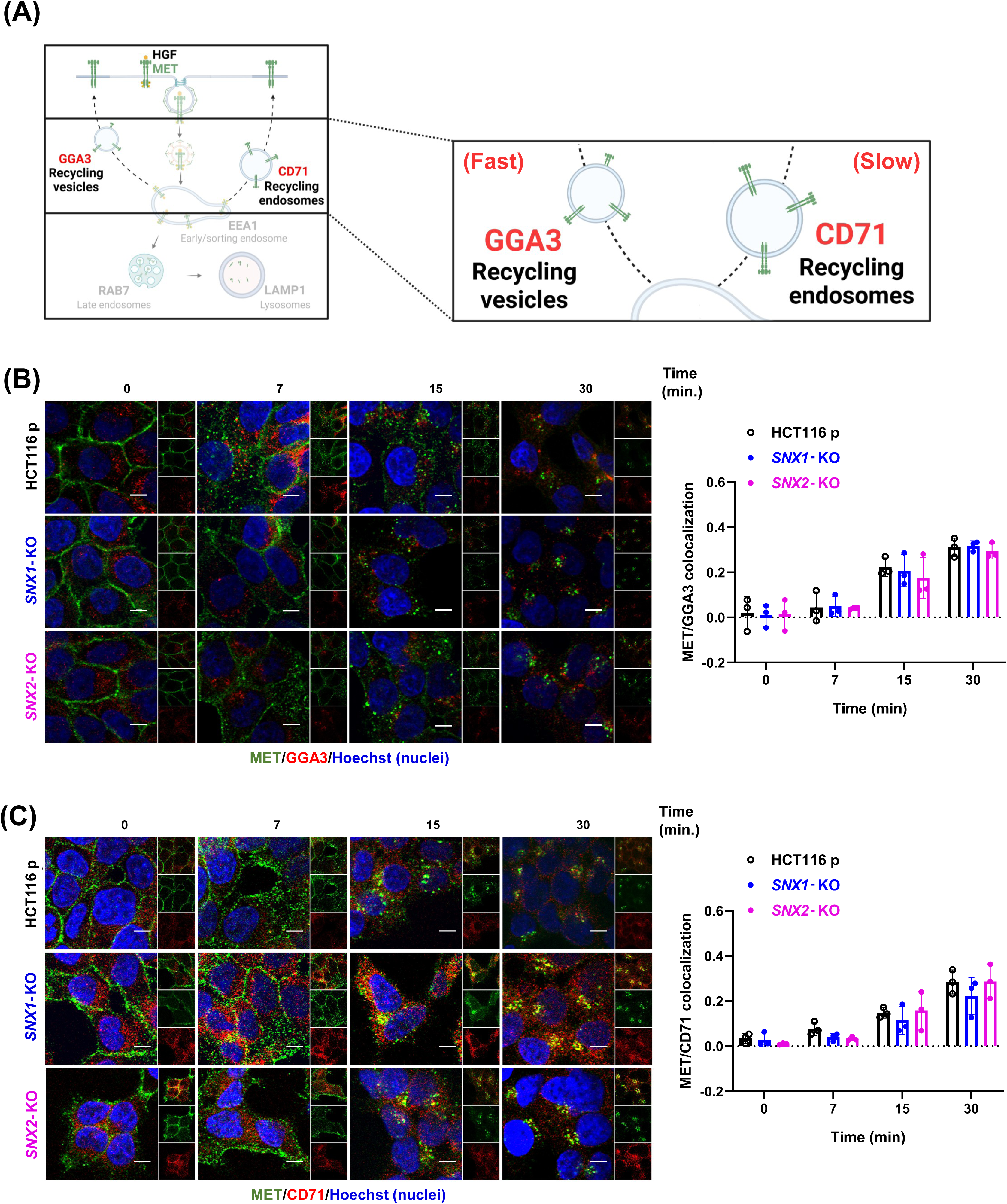
Neither *SNX1* nor *SNX2* loss alters MET flux through fast or slow endosomal recycling pathways. **A)** Schematic representation of MET trafficking to GGA3-positive fast and CD71-positive slow recycling endosomes. **B)**, **C)** Representative confocal immunofluorescence micrographs of the co-staining of MET and GGA3 **(B)** or CD71 **(C)** in parental and single *SNX1-*or *SNX2-*KO HCT116 cells. Prior to stimulation with HGF (40 ng/mL) for the indicated periods, surface MET was labeled with the anti-MET L6E7 antibody (green). Fast and slow recycling endosomes were labeled for GGA3 and CD71 (red), respectively, and nuclei were counterstained with Hoechst (blue). Scale 10 μm; 60X objective. Quantification of colocalization was assessed using the PCC (see MATERIAL AND METHODS). Data represent the mean ± SD (*N*=3).

Collectively, these data suggest that although loss of SNX2 selectively delays the early MET endosomal trafficking, its subsequent trafficking to recycling and lysosomal compartments is not detectably altered over the time course examined.

### Loss of SNX2, But Not SNX1, Enhances HGF-Induced AKT Phosphorylation

Our previous work demonstrated that acute HGF stimulation in *SNX1/2*-dKO HCT116 cells leads to hyperactivation of both MET and downstream AKT, characterized by a transient peak in pMET followed by a sustained surge in pAKT [20]. Here, we evaluated whether individual loss of *SNX1* or *SNX2* recapitulates this hyperactivation phenotype.

Following HGF stimulation of serum-starved cells, MET phosphorylation was rapidly induced within 7 min in parental and both single-KO cell lines (Fig. 7A). Although *SNX2*-KO cells displayed a trend toward sustained MET phosphorylation at later time points (30 and 120 min), overall pMET/MET ratios in both *SNX1-*KO and *SNX2-*KO cells remained comparable to parental controls throughout the time course (Fig. 7B,C).

**Figure 7:**
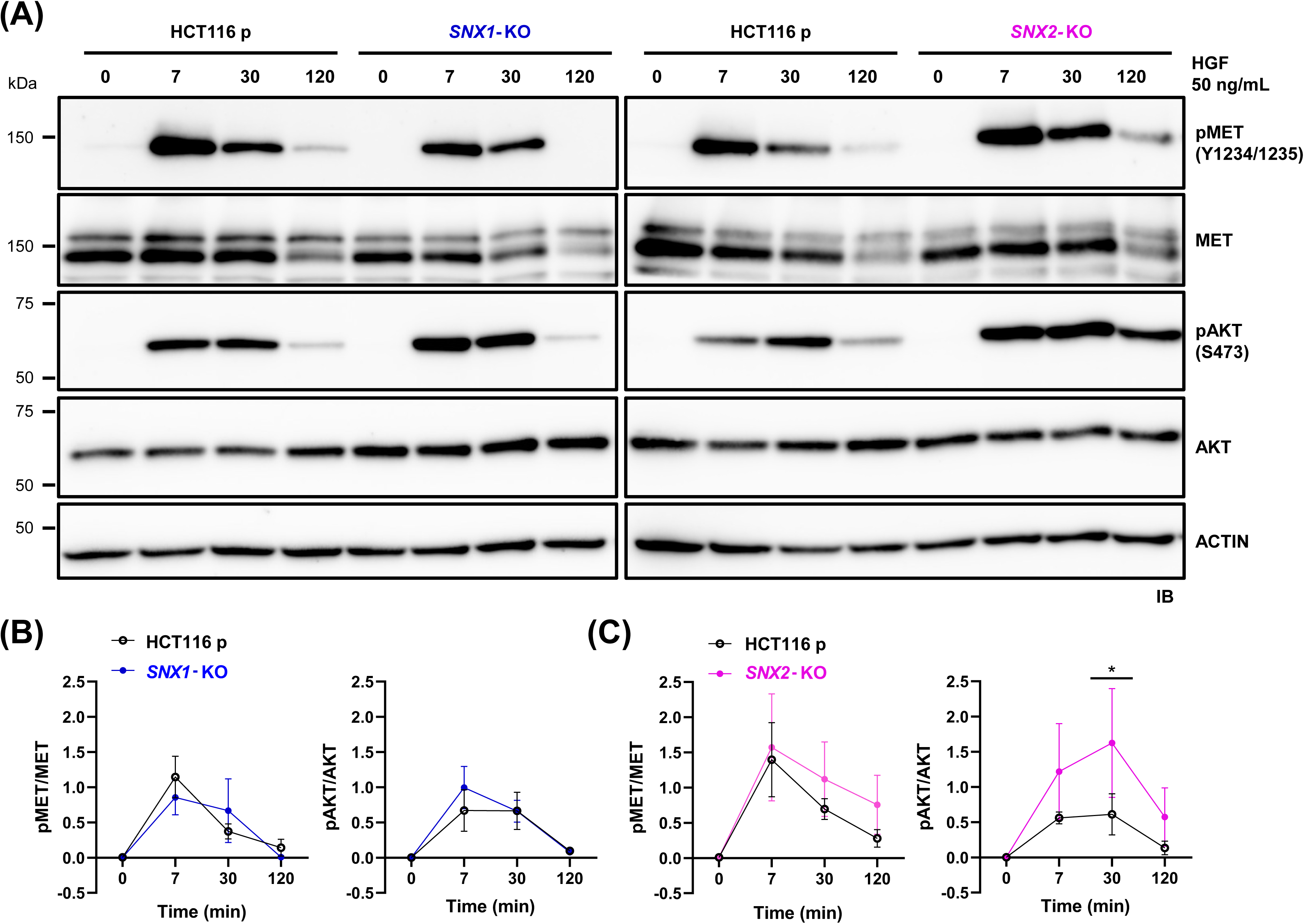
Loss of *SNX2* selectively enhances HGF-induced AKT phosphorylation downstream of MET. **A**) Serum-starved parental, *SNX1*-KO and *SNX2*-KO HCT116 cells were stimulated with 50 ng/mL HGF for the indicated periods. MET (Y1234/Y1235) and AKT (S473) phosphorylation was determined by immunoblotting, and β-Actin served as a loading control. **B, C)** Densitometric quantification of relative pMET/MET (B) and pAKT/ATK (C) ratios. Data represent the mean ± SD (*N*=3).

In contrast, downstream AKT phosphorylation was significantly increased in *SNX2*-KO cells. While parental and *SNX1*-KO cells displayed similar AKT phosphorylation kinetics, with a modest increase in pAKT/AKT ratios that plateaued between 7 and 30 min, *SNX2*-KO cells showed elevated AKT phosphorylation as early as 7 min, peaking at 30 min post-stimulation, and remaining elevated at 120 min (Fig. 7A, C). Quantification of total MET and AKT protein levels normalized to actin revealed no statistically significant differences across cell lines or time points during acute stimulation (Fig. S2), indicating that the observed signaling differences reflect changes in phosphorylation rather than protein abundance.

Collectively, these data indicate that the altered endosomal trafficking observed in *SNX2*-KO cells is associated with enhanced downstream AKT signaling, without significant changes in total MET abundance or MET phosphorylation under these acute serum-starved conditions. Furthermore, whereas dual loss of SNX1 and SNX2 produces robust MET hyperphosphorylation [20], the present data show that loss of SNX2 alone is sufficient to drive sustained downstream AKT activation following HGF stimulation.

## DISCUSSION

In this study, we dissected the role of individual SNX1 and SNX2 in modulating MET receptor trafficking and signaling. Our findings demonstrate that loss of SNX2, but not SNX1, transiently delays MET early endosomal trafficking and enhances downstream AKT activation following HGF stimulation. These findings recapitulate key features of our published observations in *SNX1/2*-dKO HCT116 cells [20] while shedding light on a functional divergence between SNX1 and SNX2. The distinct phenotypes observed in single *SNX1-*and *SNX2*-KO cells suggest that these closely related proteins possess individual functions beyond their reported redundancy in other cellular processes, such as retromer-mediated endosomal sorting [26].

Our comparative analysis of individual *SNX1*-and *SNX2*-KO HCT116 cell populations first revealed a marked reduction in SNX5 and SNX6 protein expression selectively in *SNX2*-KO cells, whereas their mRNA levels remained unaffected. Re-expression of either SNX1 or SNX2 in *SNX1/2*-dKO cells restored SNX5 and SNX6 protein levels. Notably, SNX2 was restored to levels comparable to those observed in parental cells, whereas SNX1 was expressed at substantially higher levels, suggesting that SNX1 retains the capacity to support SNX5 and SNX6 expression when present at sufficient abundance.

Together with previous reports [11,22], our findings support a post-translational co-regulation of SNX1/2/5/6 proteins, which assemble into heterodimers that function within higher-order complexes such as the ESCPE-1 [11]. Our data further suggest that SNX2 plays a dominant role in stabilizing these heterodimers in HCT116 cells, consistent with the dependence of these proteins on heterodimeric assembly [7,22]. Consequently, phenotypes observed in *SNX2*-KO cells should be interpreted as reflecting the combined loss of SNX2 and reduced SNX5 and SNX6 function. In contrast, *SNX1* loss did not destabilize SNX5 or SNX6 proteins to the same extent, although re-expression of SNX1 at sufficiently high levels restored their protein levels, further revealing a functional asymmetry between *SNX1* and *SNX2*.

The asymmetric consequences of SNX1 and SNX2 deficiency are consistent with previous genetic studies demonstrating that, while individual *Snx1*-or *Snx2*-KO mice are viable, *Snx1/2*-dKO mice exhibit embryonic lethality, underscoring a substantial functional redundancy during development [27]. However, mice retaining only a single functional allele of either *Snx1* or *Snx2* displayed distinct phenotypic outcomes. Specifically, *Snx1*^+/-^; *Snx2*^-/-^ mice were born at sub-Mendelian ratios and exhibited severe growth retardation relative to their *Snx*1^-/-^; *Snx*2^+/-^ littermates [27]. These *in vivo* genetic data demonstrate that SNX1 and SNX2 are not entirely functionally equivalent and, together with our findings, support a model in which they fulfill overlapping essential functions while making quantitatively distinct contributions within specific cellular contexts.

We observed that *SNX2* deletion increased total MET protein levels under FBS-supplemented culture conditions, whereas serum starvation restored MET expression to levels comparable to parental cells. Quantitative RT-PCR revealed equivalent *MET* transcript abundance across all cell lines, while no genotype-dependent differences in cell-surface MET were detected by FACS under either culture condition. The serum dependence of this phenotype suggests that SNX2 becomes particularly important when receptor trafficking is continuously engaged by growth-factor signaling. Serum contains multiple growth factors [28] that can trigger basal RTK activation, endocytosis, endosomal sorting, and lysosomal degradation [29]. MET can also undergo ligand-independent transactivation through crosstalk with other cell-surface receptors [30]. In HCT116 cells, the potential for this type of crosstalk is supported by the observation that both EGF (epidermal growth factor) and IGF-1 (insulin-like growth factor 1) can induce MET phosphorylation [31]. Thus, under serum-containing conditions, MET trafficking could be engaged by low-level HGF-driven MET internalization and/or heterologous cross-activation by other serum-responsive receptors, and disruption of this ongoing receptor trafficking in *SNX2*-KO cells may contribute to MET accumulation. The relative contribution of these mechanisms remains to be determined. However, the absence of increased cell-surface MET suggests that the additional receptor resides predominantly intracellularly, although the identity and functional state of this pool remain unknown.

The role of SNX1 and SNX2 in orchestrating endocytic routing of activated MET is further underscored by our spatiotemporal immunofluorescence analyses. By tracking receptor distribution across defined organellar compartments over time, we found that *SNX2* deficiency transiently delays MET trafficking to both EEA1^+^ early and RAB7^+^ late endosomes. Importantly, this occurs without detectable alterations in subsequent trafficking kinetics of MET recycling or lysosomal compartments. While these phenotypic changes partially mirror those we documented in *SNX1/2*-dKO cells [20], the decreased MET localization within RAB7^+^ endosomes was not observed in dKO cells and is specific to the single *SNX2*-KO context.

The transient delay in MET reaching RAB7^+^ endosomes is intriguing given that SNX2 deficiency is a shared genetic feature of both single *SNX2*-KO and *SNX1/2*-dKO models. This *SNX2*-KO cell-specific phenotype may primarily reflect propagation of the earlier trafficking defect, with delayed progression through early endosomes consequently postponing receptor delivery to the late endosomes. However, a critical biochemical divergence between these cell models lies in the relative abundance of the ESCPE-1 partner proteins, SNX5 and SNX6. Whereas expression of these SNXs is almost abolished in *SNX1/2*-dKO cells [20] due to post-transcriptional destabilization, a residual pool persists following single deletion of *SNX2*. As integral constituents of a higher-order complex alongside SNX1 or SNX2, SNX5 and SNX6 may modulate MET endosomal sorting dynamics. Although such a role in regulating MET trafficking remains to be formally established, previous studies have demonstrated that SNX5 and SNX6 contribute to the endosomal routing and signaling kinetics of other RTKs, including the IGF-1R and the EGFR [14,32].

Because SNX2 predominantly associates with early endosomal membranes [33], the mechanism by which its loss alters the earliest detectable steps of MET trafficking remains unresolved. At 7 min after HGF stimulation, a greater proportion of MET remained at or near the plasma membrane in *SNX2*-KO cells, concomitant with reduced localization to EEA1^+^ endosomes. This distribution suggests that the trafficking defect occurs before or during MET entry into the EEA1^+^ compartment. The phenotype could reflect delayed receptor internalization, altered routing through an EEA1^-^ intermediate compartment, or delayed maturation into EEA1^+^ endosomes following internalization. Distinguishing among these possibilities will require direct analysis of the earliest stages of MET internalization. Nonetheless, SNX2 has been reported to interact with MET independently of the receptor’s activation status [34], raising the possibility that a MET-SNX2 complex exists at or near the plasma membrane before HGF engagement, thereby positioning SNX2 to influence initial receptor internalization and early sorting dynamics.

In *SNX2*-KO cells, delayed early MET trafficking following HGF stimulation was accompanied by enhanced downstream AKT activation, consistent with observations in *SNX1/2*-dKO cells [20], whereas primary MET phosphorylation remained unaffected. These findings indicate that loss of *SNX2* alone is sufficient to alter downstream MET signaling outputs without substantially increasing receptor phosphorylation. Endocytic trafficking alterations that extend or shift the residence time of activated MET within specific intracellular compartments can dictate its spatial exposure to key scaffolding adaptors and downstream effectors [3,35]. MET activates the PI3K-AKT pathway primarily through engagement of the GAB1 (GRB2-associated binding protein 1) scaffold. GAB1 directly associates with MET or is recruited indirectly through GRB2-containing adaptor complexes, including those assembled via SHC proteins, and subsequently recruits the p85 regulatory subunit of PI3K [36–38]. Thus, delayed progression of MET from the cell surface into the early endosomal pathway could alter the duration or subcellular context in which MET engages these signaling complexes, thereby favoring sustained AKT activation. Determining where this enhanced signaling originates will require compartment-resolved analysis of MET–AKT signaling.

In conclusion, our findings challenge the classical view of SNX1 and SNX2 as largely redundant paralogs within endosomal sorting machinery and identify SNX2 as a predominant regulator of early MET endosomal trafficking in HCT116 cells. Loss of SNX2 delays receptor progression through the early endosomal pathway and sustains downstream AKT activation without substantially increasing MET phosphorylation, supporting a role for SNX2 in the spatial control of MET signaling through receptor trafficking. These findings also provide functional context for our earlier observation that SNX2 is a caspase substrate, as caspase-mediated cleavage at Asp84 disrupts its interaction with VPS35, while SNX2 depletion potentiates MET signaling [19]. Thus, proteolytic disruption of SNX2 could similarly compromise its ability to regulate MET trafficking and signaling. This possibility is particularly relevant in CRC, where SNX2 expression is reduced from early disease stages and low SNX2 expression, unlike low SNX1, is associated with poorer patient survival [19]. Important directions for future investigation include determining whether this SNX2-dependent regulatory function involves direct effects of SNX2 loss and/or the accompanying reduction in SNX5 and SNX6, whether it extends to other RTKs, and whether its disruption contributes more broadly to oncogenic signaling in CRC.

## MATERIALS AND METHODS

### Antibodies and Reagents

**Supplementary Table S1.** includes the antibodies used in this study. Recombinant human soluble TRAIL (ALX-201-073) was purchased from Enzo Life Sciences (Farmingdale). Human HGF (100-39) was from PeproTech (Cranbury). General chemicals were from Sigma-Aldrich (Burlington) and Thermo Fisher Scientific (Waltham).

### Cell Culture, Transfection, and Treatments

The human colorectal carcinoma cell line HCT116 (CCL-247) was purchased from ATCC (Cedarlane), and human embryonic kidney 293T (HEK293T; CRL-3216) cells were kindly provided by Dr. F.-P. Gendron (Université de Sherbrooke, Sherbrooke, QC, CANADA). Cells were maintained in Dulbecco’s modified Eagle’s medium (DMEM) supplemented with 10% fetal bovine serum (FBS), 2 mM L-glutamine, and penicillin-streptomycin antibiotics (Wisent Inc.) in a 5% CO_2_ atmosphere at 37°C.

The *SNX1* or *SNX2* single KO cells were generated using CRISPR/Cas9 gene-editing technology. The U6-gRNA/CMV-eCas9-2a-tGFP vector (Sigma-Aldrich) containing the guide RNA targeting *SNX1* (5’-GGACAACACGGCATTGTCA-3’) or *SNX2* (5’-AAACGATTCGAAAAGAAGT-3’) was transfected into HCT116 cells using Lipofectamine 2000 (Thermo Fisher Scientific), following the manufacturer’s instructions. After 48 h, GFP-positive cells were individually sorted into 96-well plates (BD FACSAria™ III; BD Biosciences). *SNX1/2*-dKO cells were generated as described elsewhere [20]. Clones were screened for loss of *SNX1* or *SNX2* expression by immunoblotting and genomic DNA sequencing, using the following primer pairs: SNX1: 5’-GGCTTTGGTTCTCCAGGGAT-3’; 5’-AAGGCTCTGGTGACAAGCTC-3’; SNX2: 5’-GAGCAAATCTGTGGAATATTAAAGG-3’; 5’-AAACTGGAAGGAAGAAATCCT-3’. For signaling experiments, serum-starved cells were stimulated with HGF (50 ng/mL) for the indicated times.

### Retrovirus Production and Transduction

*SNX1* and *SNX2* coding sequences were subcloned via EcoRI and NotI digestion into the retroviral vector pLPCX to stably re-express *SNX1* or *SNX2* in dKO cells. HEK293T cells were transfected with pLPCX-SNX1, pLPCX-SNX2, or the empty pLPCX vector along with the pVPack-GP and pVPack-VSV-G vectors (Agilent), using Lipofectamine 2000 (Invitrogen). The supernatant was collected and filtered after 48 h and added to *SNX1/2*-dKO HCT116 cells with 4 µg/mL of polybrene (Sigma-Aldrich). Cells were cultured for 48 h before selection with 1 µg/mL of puromycin (Sigma-Aldrich). Populations of *SNX1/2*-dKO cells re-expressing SNX1 or SNX2 were expanded from puromycin-resistant colonies.

### Immunoblotting

Cells were washed with ice-cold PBS (123 mM NaCl, 10.4 mM Na₂HPO₄, 3 mM KH₂PO₄, pH 7.4) and lysed for 30 min on ice in Triton X-100 lysis buffer (50 mM HEPES, pH 7.4, 150 mM NaCl, 0.5% (v/v) Triton X-100, 10% (v/v) glycerol, 2 mM EGTA, and 1.5 mM MgCl₂) supplemented with protease inhibitors (1 mM 1,10-orthophenanthroline, 1 mM Na₃VO₄, 1 mM NaF, 50 μM 3,4-dichloroisocoumarin, 10 μM E64, and 10 μM leupeptin). Lysates were clarified by centrifugation at 18,000 × g for 15 min, and protein concentration was determined using the bicinchoninic acid (BCA) assay (Thermo Fisher Scientific). Equivalent protein amounts were separated by 2-Amino-2-methyl-1,3-propanediol (ammediol) SDS-PAGE and transferred onto PVDF membranes (EMD Millipore) using transfer buffer (10 mM N-cyclohexyl-3-aminopropanesulfonic acid, pH 11, 10% (v/v) methanol). Membranes were blocked in PBS containing 0.1% (v/v) Tween-20 and either 5% (w/v) non-fat dry milk (Carnation) or 3% (w/v) BSA (Sigma-Aldrich) for phosphoprotein detection. Blots were incubated overnight at 4°C with primary antibodies (Table S1), followed by a 1-h incubation with HRP-conjugated secondary antibodies. Signals were detected using Immobilon Crescendo Western HRP substrate (EMD Millipore) or Clarity Max Western ECL Substrate (Bio-Rad). Images were acquired using a ChemiDoc MP Imaging System (Bio-Rad), and densitometric analyses were performed using Image Lab 6.1 (Bio-Rad). When required, membranes were stripped using a buffer containing 62.5 mM Tris, pH 6.8, 2% SDS, and 100 mM β-mercaptoethanol at 50°C for 30 min under gentle agitation, then re-probed with different primary antibodies.

### Immunofluorescence

Cells were cultured on poly-L-lysine-coated (Sigma-Aldrich) glass coverslips (Thermo Fisher Scientific) and allowed to adhere for 48 h. Before stimulation, cells were serum-starved for 1 h in DMEM containing 0.2% (w/v) BSA and 25 mM HEPES. To label surface MET, cells were incubated for 1 h at 4 °C with the anti-MET L6E7 antibody, then stimulated with HGF. Following stimulation, cells were fixed with 3% (w/v) paraformaldehyde (Electron Microscopy Sciences) for 30 min, permeabilized for 10 min in 0.1% (v/v) Triton X-100, and blocked for 30 min with 10% (v/v) goat serum. Cells were subsequently incubated for 1 h with specific primary antibodies, followed by 45 min of incubation with Alexa Fluor-conjugated secondary antibodies. Coverslips were mounted onto glass slides (Globe Scientific Inc.) using ProLong Glass Antifade Mountant (Thermo Fisher Scientific). Images were acquired with an Olympus FV1000 Spectral SIM confocal microscope operated with FluoView FV-10 ASW software v4.2. Images were analyzed using CellProfiler v4.0.7 [39] using Pearson’s overlap to quantify colocalization of MET with the different subcellular compartment markers.

### Flow Cytometry

The surface levels of MET were analyzed by flow cytometry. Cells were seeded in 35-mm dishes and collected after 48 h using cold flow cytometry buffer (PBS supplemented with 2% (v/v) FBS). Cells were incubated with an Alexa Fluor 488-conjugated anti-MET antibody diluted in the same buffer. To validate the assay, HCT116 cells transfected with *MET*-targeting DsiRNAs (Integrated DNA Technologies) were used as a control for reduced MET expression. Data were acquired using a Cytek Guava EasyCyte flow cytometer (Cytek Biosciences Inc.) and analyzed using Kaluza Analysis v2.4 (Beckman Coulter).

### Gene Expression

The levels of *MET* mRNA were determined by qRT-PCR at our institutional *Plateforme de RNomique* using the following primer pairs: 5’-TGGCTACACACTGGTTATCACTGG-3’ and 5’-ACTGGAAATGTCTGCAGCCCAA-3’; *MRPL19* (mitochondrial ribosomal protein L19), *PUM1* (pumilio RNA-binding family member 1), and *YWHAZ* (tyrosine 3-monooxygenase/tryptophan 5-monooxygenase activation protein zeta) were used as reference genes. Messenger RNA was isolated from cells by TRIzol reagent (ThermoFisher Scientific) followed by chloroform extraction and purification with RNeasy Mini spin columns (QIAGEN). First-strand cDNA was synthesized from 1 μg of total RNA using Transcriptor Reverse Transcriptase (Roche) according to the manufacturer’s instructions. Quantitative PCR was performed on a CFX Opus 384 Real-Time PCR System (Bio-Rad) using 3.33 ng of cDNA, 2× SYBR Green Supermix (Bio-Rad), and gene-specific primer pairs. Relative gene expression was quantified using the qBASE framework [40] and normalized to *MRPL19*, *PUM1*, and *YWHAZ* expression.

### Statistical Analyses

Data were analyzed using the GraphPad Prism software (v.9.5.1). For each of three independent experiments, microscopy images were acquired from three different fields containing approximately 60 cells per condition. Comparisons among parental, *SNX1-*, and *SNX2*-KO HCT116 cells were performed using one-way or two-way ANOVA, as appropriate. Two-way ANOVA was applied along with Sidak’s multiple-comparison correction. Statistical significance was indicated as follows: *, *p* ≤ 0.05; **, *p* ≤ 0.01; ***, *p* ≤ 0.001.

## Supporting information

https://usherbrooke.sharepoint.com/:b:/r/sites/DLab/Documents%20partages/General/Laiyen%20Garcia%20Delgado/PAPER/Single%20KO/BioRxiv/Garcia%20et%20al,

## DATA AVAILABILITY

All data are included in the manuscript and/or supplemental file.

## ACKNOWLEDGEMENTS

This work was supported by the Canadian Institutes for Health Research (CIHR) PJT-162354 (to J-BD, CS, and CLL). LGD was supported by an IRCUS excellence and a FRQ studentships and AD by a CIHR studentship award. J-BD, CS, and CLL are members of the FRQS-funded Centre de Recherche du CHUS.

## ETHICS STATEMENT

The authors have nothing to report.

## AUTHORS CONTRIBUTION

**LGD** designed the experiments, performed the research, analyzed the data, and wrote the manuscript.

**AD** designed the experiments, performed the research, and analyzed the data.

**CLL** designed the experiments, analyzed the data, and wrote the manuscript.

**CS** designed the experiments, analyzed the data, and wrote the manuscript.

**J-BD** designed the experiments, analyzed the data, and wrote the manuscript

## REFERENCES

1. Zastrow, M. von & Sorkin, A. Mechanisms for Regulating and Organizing Receptor Signaling by Endocytosis. Annu. Rev. Biochem. 90, 709–737 (2021).

2. Bergeron, J. J. M., Di Guglielmo, G. M., Dahan, S., Dominguez, M. & Posner, B. I. Spatial and Temporal Regulation of Receptor Tyrosine Kinase Activation and Intracellular Signal Transduction. Annu. Rev. Biochem. 85, 573–597 (2016).

3. Joffre, C. et al. A direct role for Met endocytosis in tumorigenesis. Nat. Cell Biol. 13, 827– 837 (2011).

4. Pinilla-Macua, I., Surve, S. & Sorkin, A. Cell migration signaling through the EGFR-VAV2-Rac1 pathway is sustained in endosomes. J. Cell Sci. 138, jcs263541 (2025).

5. Schultz, D. F., Billadeau, D. D. & Jois, S. D. EGFR trafficking: effect of dimerization, dynamics, and mutation. Front. Oncol. 13, 1258371 (2023).

6. Teasdale, R., Loci, D., Houghton, F., Karlsson, L. & Gleeson, P. A. A large family of endosome-localized proteins related to sorting nexin 1. Biochem. J. 358, 7–16 (2001).

7. Wassmer, T. et al. A loss-of-function screen reveals SNX5 and SNX6 as potential components of the mammalian retromer. J. Cell Sci. 120, 45–54 (2007).

8. Klinger, S. C., Siupka, P. & Nielsen, M. S. Retromer-Mediated Trafficking of Transmembrane Receptors and Transporters. Membranes 5, 288–306 (2015).

9. Cozier, G. E. et al. The Phox Homology (PX) Domain-dependent, 3-Phosphoinositide-mediated Association of Sorting Nexin-1 with an Early Sorting Endosomal Compartment Is Required for Its Ability to Regulate Epidermal Growth Factor Receptor Degradation *. J. Biol. Chem. 277, 48730–48736 (2002).

10. Wang, Y., Zhou, Y., Szabo, K., Haft, C. R. & Trejo, J. Down-Regulation of Protease-activated Receptor-1 Is Regulated by Sorting Nexin 1. Mol. Biol. Cell 13, 1965–1976 (2002).

11. Kvainickas, A. et al. Cargo-selective SNX-BAR proteins mediate retromer trimer independent retrograde transport. J. Cell Biol. 216, 3677–3693 (2017).

12. Kurten, R. C., Cadena, D. L. & Gill, G. N. Enhanced Degradation of EGF Receptors by a Sorting Nexin, SNX1. Science 272, 1008–1010 (1996).

13. Gullapalli, A. et al. A Role for Sorting Nexin 2 in Epidermal Growth Factor Receptor Down-regulation: Evidence for Distinct Functions of Sorting Nexin 1 and 2 in Protein Trafficking. Mol. Biol. Cell 15, 2143–2155 (2004).

14. Zhou, Q. et al. Upregulation of SNX5 predicts poor prognosis and promotes hepatocellular carcinoma progression by modulating the EGFR-ERK1/2 signaling pathway. Oncogene 39, 2140–2155 (2020).

15. Hu, P. et al. SNX6 predicts poor prognosis and contributes to the metastasis of pancreatic cancer cells via activating epithelial-mesenchymal transition. Acta Biochim. Biophys. Sin. 50, 1075–1084 (2018).

16. Atwell, B., Chen, C.-Y., Christofferson, M., Montfort, W. R. & Schroeder, J. Sorting nexin-dependent therapeutic targeting of oncogenic epidermal growth factor receptor. Cancer Gene Ther. 30, 267–276 (2023).

17. Nishimura, Y., Takiguchi, S., Ito, S. & Itoh, K. Evidence that depletion of the sorting nexin 1 by siRNA promotes HGF-induced MET endocytosis and MET phosphorylation in a gefitinib-resistant human lung cancer cell line. Int. J. Oncol. 44, 412–426 (2014).

18. Ogi, S. et al. Sorting nexin 2-mediated membrane trafficking of c-Met contributes to sensitivity of molecular-targeted drugs. Cancer Sci. 104, 573–583 (2013).

19. Duclos, C. M. et al. Caspase-mediated proteolysis of the sorting nexin 2 disrupts retromer assembly and potentiates Met/hepatocyte growth factor receptor signaling. Cell Death Discov. 3, 1–12 (2017).

20. Garcia Delgado, L., et al. Spatiotemporal regulation of the hepatocyte growth factor receptor MET activity by sorting nexins 1/2 in HCT116 colorectal cancer cells. Biosci. Rep. BSR20240182 (2024) doi:10.1042/BSR20240182.

21. Uhlén, M. et al. Tissue-based map of the human proteome. Science 347, 1260419 (2015).

22. Simonetti, B., Danson, C. M., Heesom, K. J. & Cullen, P. J. Sequence-dependent cargo recognition by SNX-BARs mediates retromer-independent transport of CI-MPR. J. Cell Biol. 216, 3695–3712 (2017).

23. Adler, J. & Parmryd, I. Quantifying colocalization by correlation: The Pearson correlation coefficient is superior to the Mander’s overlap coefficient. Cytometry A **77A**, 733–742 (2010).

24. Dunn, K. W., Kamocka, M. M. & McDonald, J. H. A practical guide to evaluating colocalization in biological microscopy. Am. J. Physiol. - Cell Physiol. 300, C723–C742 (2011).

25. Hammond, D. E., Urbé, S., Vande Woude, G. F. & Clague, M. J. Down-regulation of MET, the receptor for hepatocyte growth factor. Oncogene 20, 2761–2770 (2001).

26. Rojas, R., Kametaka, S., Haft, C. R. & Bonifacino, J. S. Interchangeable but essential functions of SNX1 and SNX2 in the association of retromer with endosomes and the trafficking of mannose 6-phosphate receptors. Mol. Cell. Biol. 27, 1112–1124 (2007).

27. Schwarz, D. G., Griffin, C. T., Schneider, E. A., Yee, D. & Magnuson, T. Genetic Analysis of Sorting Nexins 1 and 2 Reveals a Redundant and Essential Function in Mice. Mol. Biol. Cell 13, 3588–3600 (2002).

28. Wijerathna-Yapa, A. & Sokolenko, S. What Is in Fetal Bovine Serum Used for Cell Culture? Integrated Proteomics and Peptidomics Reveal Grade-Dependent Molecular Profiles. J. Proteome Res. 25, 2772–2788 (2026).

29. Goh, L. K. & Sorkin, A. Endocytosis of Receptor Tyrosine Kinases. Cold Spring Harb. Perspect. Biol. 5, a017459 (2013).

30. Viticchiè, G. & Muller, P. A. J. c-Met and Other Cell Surface Molecules: Interaction, Activation and Functional Consequences. Biomedicines 3, 46–70 (2015).

31. Song, N. et al. Dual inhibition of MET and SRC kinase activity as a combined targeting strategy for colon cancer. Exp. Ther. Med. 14, 1357–1366 (2017).

32. Bareja, A., Hodgkinson, C. P., Soderblom, E., Waitt, G. & Dzau, V. J. The proximity-labeling technique BioID identifies sorting nexin 6 as a member of the insulin-like growth factor 1 (IGF1)–IGF1 receptor pathway. J. Biol. Chem. 293, 6449–6459 (2018).

33. Teasdale, R. D. & Collins, B. M. Insights into the PX (phox-homology) domain and SNX (sorting nexin) protein families: structures, functions and roles in disease. Biochem. J. 441, 39–59 (2011).

34. Schaaf, C. P. et al. Novel interaction partners of the TPR/MET tyrosine kinase. FASEB J. Off. Publ. Fed. Am. Soc. Exp. Biol. 19, 267–269 (2005).

35. Barrow-McGee, R. & Kermorgant, S. Met endosomal signalling: in the right place, at the right time. Int. J. Biochem. Cell Biol. 49, 69–74 (2014).

36. Ponzetto, C. et al. A Novel Recognition Motif for Phosphatidylinositol 3-Kinase Binding Mediates Its Association with the Hepatocyte Growth Factor/Scatter Factor Receptor. Mol. Cell. Biol. 13, 4600–4608 (1993).

37. Fixman, E. D., Fournier, T. M., Kamikura, D. M., Naujokas, M. A. & Park, M. Pathways downstream of Shc and Grb2 are required for cell transformation by the tpr-Met oncoprotein. J. Biol. Chem. 271, 13116–13122 (1996).

38. Holgado-Madruga, M., Moscatello, D. K., Emlet, D. R., Dieterich, R. & Wong, A. J. Grb2-associated binder-1 mediates phosphatidylinositol 3-kinase activation and the promotion of cell survival by nerve growth factor. Proc. Natl. Acad. Sci. U. S. A. 94, 12419–12424 (1997).

39. Stirling, D. R. et al. CellProfiler 4: improvements in speed, utility and usability. BMC Bioinformatics 22, 433 (2021).

40. Hellemans, J., Mortier, G., De Paepe, A., Speleman, F. & Vandesompele, J. qBase relative quantification framework and software for management and automated analysis of real-time quantitative PCR data. Genome Biol. 8, R19 (2007).

