## Supplementary material for "SNX2, but not SNX1, regulates the hepatocyte growth factor receptor MET in HCT116 colorectal cancer cells": https://usherbrooke.sharepoint.com/:b:/r/sites/DLab/Documents%20partages/General/Laiyen%20Garcia%20Delgado/PAPER/Single%20KO/BioRxiv/Garcia%20et%20al,

**Title:**

**Supplementary Material:**

Supplementary Table (Table S1)
**Table S1:** List of antibodies used in the study.

Supplementary Figures (Figures S1-S2)
**Figure S1:** Validation of CRISPR/Cas9-mediated knockout of *SNX1* and *SNX2* in HCT116 cells.
**Figure S2:** Total MET and AKT levels remain unchanged between parental and single KO cells upon HGF stimulation.

**Table S1. Antibodies used.**

| Antibodies for immunoblotting | | | |
|---|---|---|---|
| **Antibody** | **Dilution** | **Catalog number** | **Source** |
| MET (L41G3) | 1:2000 | 3148 | Cell Signaling Technology (Danvers, MA, USA) |
| SNX5 (F-11) | 1:1000 | sc-515215 | Santa Cruz Biotechnology (Dallas, TX, USA) |
| SNX6 (D-5) | 1:1000 | sc-365965 | Santa Cruz Biotechnology |
| SNX2 | 1:1000 | Y00322-002 | Immune Biosolutions (Sherbrooke, QC, Canada) |
| SNX1 (51) | 0.25 µg/mL | 611482 | BD Biosciences (Franklin Lakes, NJ, USA) |
| Phospho-MET (Tyr1234/1235) (D26) | 1:1000 | 3077 | Cell Signaling Technology |
| Phospho-AKT (Ser473) | 1:500 | 9271 | Cell Signaling Technology |
| AKT | 1:1000 | 9272 | Cell Signaling Technology |
| Actin (AC-40) | 1:5000 | A3853 | Sigma-Aldrich (St-Louis, MO, USA) |
| HSP90 (68) | 1:10000 | 610419 | BD Biosciences |
| Anti-mouse IgG, HRP-linked | 1:5000 | 7076 | Cell Signaling Technology |
| Anti-rabbit IgG, HRP-linked | 1:5000 | 7074 | Cell Signaling Technology |
| Anti-chicken IgG, HRP-linked | 1:5000 | A16054 | Thermo Fisher Scientific (Waltham, MA, USA) |

| Antibodies for immunofluorescence | | | |
|---|---|---|---|
| **Antibody** | **Dilution** | **Catalog number** | **Source** |
| MET (L6E7) | 1:1000 | 8741 | Cell Signaling Technology |
| EEA1 | 10 µg/mL | PA1-063A | Thermo Fisher Scientific |
| CD71 (D7G9X) | 1:100 | 13113 | Cell Signaling Technology |

| | | | |
|---|---|---|---|
| LAMP1 (D2D11) | 5 µg/mL | 9091 | Cell Signaling Technology |
| RAB7 (D95F2) | 1:50 | 9367 | Cell Signaling Technology |
| GGA3 | 1 µg/mL | PA5-82888 | Thermo Fisher Scientific |
| Goat anti-Rabbit IgG (H+L) Cross-Adsorbed Secondary Antibody, Alexa Fluor™ 405 | 12 ng/mL | A31556 | Thermo Fisher Scientific |
| Donkey anti-Rabbit IgG (H+L) Highly Cross-Adsorbed Secondary Antibody, Alexa Fluor™ 488 | 12 ng/mL | A21206 | Thermo Fisher Scientific |
| Donkey anti-Rabbit IgG (H+L) Highly Cross-Adsorbed Secondary Antibody, Alexa Fluor™ 594 | 12 ng/mL | A21207 | Thermo Fisher Scientific |
| Donkey anti-Mouse IgG (H+L) Highly Cross-Adsorbed Secondary Antibody, Alexa Fluor™ 488 | 12 ng/mL | A21202 | Thermo Fisher Scientific |
| Donkey anti-Mouse IgG (H+L) Highly Cross-Adsorbed Secondary Antibody, Alexa Fluor™ 594 | 12 ng/mL | A21203 | Thermo Fisher Scientific |

| **Antibodies for cytometry** | | | |
|---|---|---|---|
| **Antibody** | **Dilution** | **Catalog number** | **Source** |
| Human HGFR/c-MET Alexa Fluor® 488-conjugated | 5 µL/$10^6$ cells | FAB3582G | R&D systems (Minneapolis, MN, USA) |

### Figures S1

(A)

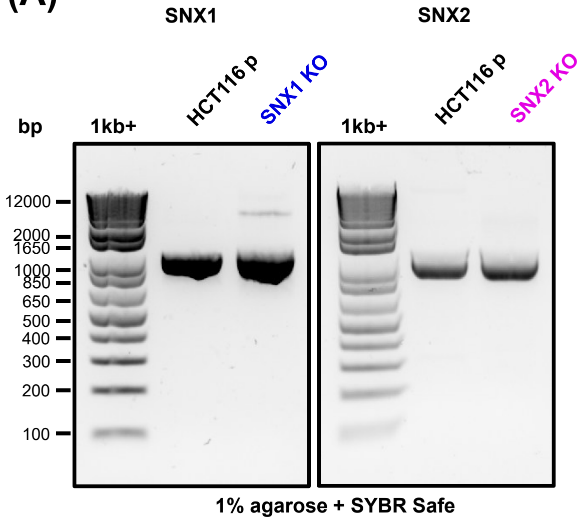


(B)

| Cells | SNX1 sequence | SNX2 sequence |
|---|---|---|
| **HCT116 p** | wt | wt |
| **SNX1 KO** | 11 nt deletion | wt |
| **SNX2 KO** | wt | 3 nt deletion<br>1 nt insertion |

**Figure S1: Validation of CRISPR/Cas9-mediated knockout of *SNX1* and *SNX2* in HCT116 cells. A** Genomic DNA from parental and single KO cells was subjected to PCR amplification of a fragment of *SNX1* or *SNX2* that includes the CRISPR/Cas9 target region. The following primer pairs were used: SNX1-F: 5'-GGCTTTGGTTCTCCAGGGAT-3'; SNX1-R: 5'-AAGGCTCTGGTGACAAGCTC-3'; SNX2-F: 5'-GAGCAAATCTGTGGAATATTAAAGG-3'; SNX2-R: 5'-AAACTGGAAGGAAGAAATCCT-3', and the amplicons were next sequenced. **B** The table summarizes the modifications in the sequence of *SNX1* and *SNX2* that result in the depletion of both proteins.

Figure S2

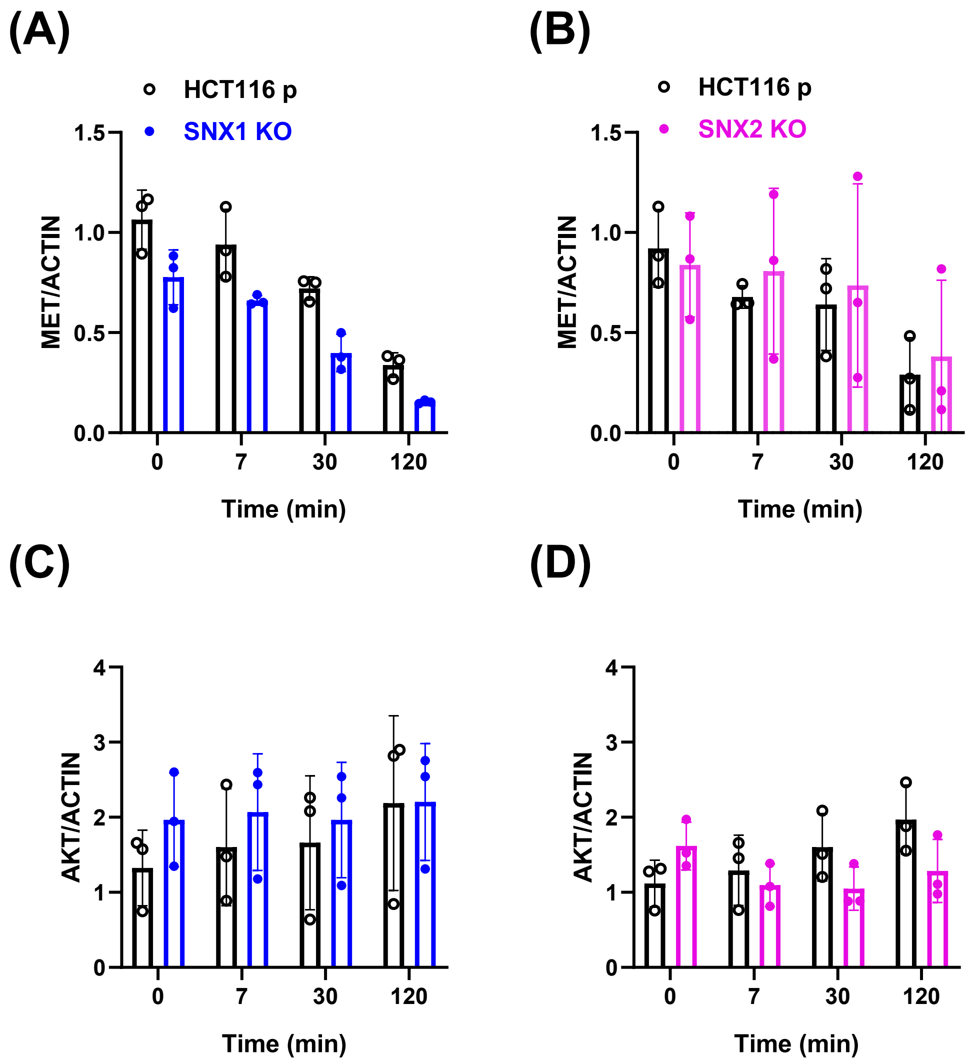


**Figure S2: Total MET and AKT levels remain unchanged between parental and single KO cells upon HGF stimulation.** Serum-starved cells were stimulated with 50 ng/mL HGF for the indicated period. Graphs show the densitometric quantification of data from Figure 7A, expressed as the ratio of total MET **(A, B)** or total AKT **(C, D),** normalized to the actin loading control. Data represent the mean ± SD (N=3).
